# Benchmarking multi-omics integration algorithms across single-cell RNA and ATAC data

**DOI:** 10.1101/2023.11.15.564963

**Authors:** Chuxi Xiao, Yixin Chen, Lei Wei, Xuegong Zhang

## Abstract

Recent advancements in single-cell sequencing technologies have generated extensive omics data in various modalities and revolutionized cell research, especially in the single cell RNA and ATAC data. The joint analysis across scRNA-seq data and scATAC-seq data has paved the way to comprehending the cellular heterogeneity and complex cellular regulatory networks. Multi-omics integration is gaining attention as an important step in joint analysis, and the number of computational tools in this field is growing rapidly. In this paper, we benchmarked 12 multi-omics integration methods on three integration tasks via qualitative visualization and quantitative metrics, considering six main aspects that matter in multi-omics data analysis. Overall, we found that different methods have their own advantages on different aspects, while some methods outperformed other methods in most aspects. We therefore provided guidelines for selecting appropriate methods for specific scenarios and tasks to help obtain meaningful insights from multi-omics data integration.

## Introduction

Over the past few years, single-cell sequencing technologies have emerged, generating massive of omics data across various modalities. Single-cell RNA-seq (scRNA-seq) and single cell ATAC-seq (scATAC-seq) data are two major modalities, providing a wealth of information about gene expression regulation. Additionally, new sequencing technologies have been developed to profile paired multi-omics features within a same cell [1–4]. Conjoint analysis of these single-cell RNA and ATAC data can greatly help obtain a more comprehensive view of gene regulation within single cells.

Integration of multi-omics data is the initial phase of the conjoint analysis, with the objective of aligning data from different omics into a unified latent space. This integration can facilitate a deeper understanding of cell-specific regulatory networks by inferring upstream regulatory factors [5], and can help identify more cell clusters and biomarkers with potential clinical relevance [6,7]. Integration analysis on single-cell RNA and ATAC data has been becoming more and more common and necessary in recent research.

Many tools have been developed for single-cell RNA and ATAC data integration. Here we classify these methods into three categories according to the data modalities they designed for: unpaired integration, paired integration and paired-guided integration [8,9]. The unpaired integration methods are designed for single-cell RNA and ATAC data derived from the same tissue but different cells [8]. In this category, UnionCom [10] and MMD-MA [11] perform alignment by matching the distance matrixes of cells or low-dimensional distributions of different omics, respectively. LIGER [12] performs integrative non-negative matrix factorization (iNMF) to learn factors that correspond to different biologically signals. BindSC [13] and Seurat v3 [14] are both based on canonical correlation analysis (CCA), and Seurat v3 additionally combines CCA with graphs and uses mutual nearest neighbors (MNNs) to identify anchors. scDART [15] defines a gene activity function module to generate scATAC-seq data as pseudo scRNA-seq data, and then integrates them into a unified latent space. scJoint [16] uses labeled scRNA-seq data and unlabeled scATAC-seq data for semi-supervised learning. GLUE [17] adopts a framework that combines omics-specific autoencoders with graph-based coupling and adversarial alignment for unpaired integration.

The paired integration methods aim to deal with multi-omics data simultaneously profiling the same cell [8]. MOFA+ [18] gains a low-dimensional representation across different omics using variational inference. scAI [7] is based on matrix factorization that learns a cell-cell similarity matrix. Seurat v4 [19] establishes a weighted graph of cell–cell relationships, and then completes the integration by manipulating the constructed graph. scMVP [20] proposes a clustering consistency-constrained multi-view variational auto-encoder model (VAE) to learn a common latent representation, and reconstructs different omics layers through different channels. TotalVI [21] uses a probabilistic latent variable model to learn a low-dimensional representation across omics and introduces different modeling strategies for different omics.

The paired-guided integration methods, also known as multiome-guided integration methods [9], are designed to use paired multi-omics data to assist the integration of unpaired data. MultiVI [22] and Cobolt [23] both establish a deep generative model for integrative analysis of scRNA-seq, scATAC-seq and paired multi-modal data, assuming different distributions for each omics. MultiVI additionally uses KL divergence loss to align integration.

With the rapid growth of these methods for integrating single-cell RNA and ATAC data, a systematic evaluation becomes essential to navigate method selection in practical experiments. Recently, Lee et al. proposed a benchmark study to compare 7 integration methods [9], but both the latest deep learning-based methods such as scMVP, scDART, scJoint etc., and paired integration methods were missed in this study. Moreover, this work placed a significant focus on the details of multi-omics data, such as sequencing depths, while not paying sufficient attention to the downstream analysis stemming from integration. Here we established a benchmark that compared 12 popular methods which covered all the three categories of integration methods, that is, paired methods (scMVP [20], MOFA+ [18]), paired-guided methods (MultiVI [22], Cobolt [23]) and unpaired methods (scDART [15], UnionCom [10], MMD-MA[11], scJoint [16], Harmony [24], Seurat v3 [14], LIGER [12], and GLUE [17]). We considered four aspects to evaluate the accuracy of integration: the extent of mixing among different omics, the cell type conservation, the single-cell level alignment accuracy, and how well the expected trajectory was preserved. We also evaluated the time scalability of these methods. Moreover, we evaluated the ease of use for each method based on our experience. According to these results, we provided guidelines for method selection in different situations.

## Results

### Overview of benchmarking strategies

We benchmarked 12 popular methods in the following three categories (Fig. 1): two popular integration methods designed for paired datasets (scMVP [20], MOFA+ [18]), two popular methods belong to the paired-guided integration category (MultiVI [22], Cobolt [23]), and eight integration methods that could be used for both paired and unpaired datasets (scDART [15], UnionCom [10], MMD-MA[11], scJoint [16], Harmony [24], Seurat v3 [14], LIGER [12], and GLUE [17]). We benchmarked these methods in three datasets to evaluate their performance in different single-cell RNA and ATAC data integration tasks (Fig. 1): a P0 mouse cerebral cortex dataset with 5,081 cells generated by droplet-based SNARE-seq [1] for paired integration (Dataset-P), and 1469 cells with an expected cell trajectory that extracted from this paired dataset for integration with a trajectory (Dataset-T), and a human uterus datasets with 8,237 cells for scRNA-seq [25] and 8,314 cells for scATAC-seq [26] for unpaired integration (Dataset-U). These three datasets each posed a unique challenge, and could stand for different integration application scenarios.

**Fig. 1.**
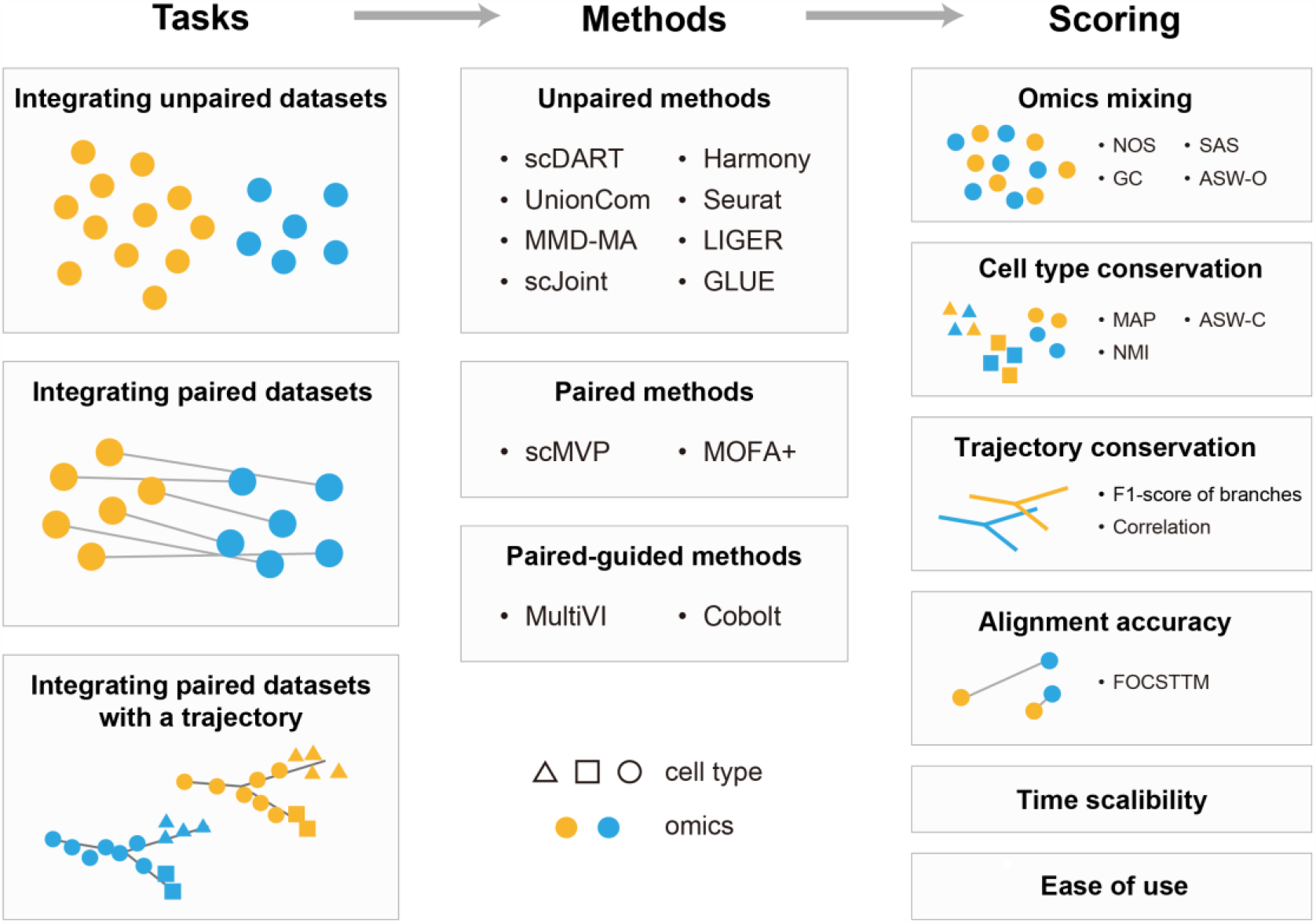
Schematic diagram of the benchmarking workflow.

We visualized the latent embedding of each method by Uniform Manifold Approximation and Projection (UMAP). We colored each cell by its omics type or cell type, respectively, to assess whether cells of the same cell type derived from different omics were clustered in the latent space. Furthermore, we carried out four kinds of metrics to evaluate the accuracy of the integration results (Fig. 1). The extent of mixing among omics (omics mixing in short) focuses on how well two omics are integrated with each other in the latent space (Fig. 1), and it was measured via the neighborhood overlap score (NOS) [15], the graph connectivity (GC) [17,27], the Seurat alignment score (SAS) [17,28], and the average silhouette width across omics (ASW-O) [17,27] (Methods). The cell type conservation evaluates whether cells of the same type are well clustered, and whether cells of different types are separated from each other (Fig. 1). It was measured by the mean average precision (MAP) [17], the average silhouette width (ASW) [17,27], and the normalized mutual information (NMI) [27] (Methods). For datasets with an expected trajectory, the trajectory conservation was measured by the F1 score of branches [15,29], and the Spearman’s and Pearson’s correlation between trajectories generated in the latent space [15] (Methods). When dealing with paired datasets, the single-cell level alignment accuracy was evaluated by FOSCTTM to assess whether the two omics data of a same cell distributed near each other in the latent space [17,30] (Methods). We also recorded the running time when testing these three datasets with different number of cells to compare the scalability of each method. Besides, we graded these methods based our experience to evaluate their ease of use.

Requirements on data preprocessing of different integration methods are inconsistent, and all methods have some adjustable parameters. We ran each integration method according to the settings mentioned in its original paper. If there are no instructions in the original papers, we ran the methods with or without scaling and highly variable gene (HVG) selection, respectively, and then chose the best results and the corresponding settings for benchmarking (Methods).

### Benchmarking results on the unpaired dataset

First, we focused on how the eight methods available for unpaired tasks performed on the unpaired dataset (Dataset-U). The four cell types (endothelial, fibroblast, macrophage and smooth muscle cell) could be divided clearly in each omics (Fig. 2a). From the UMAP colored by omics type (Fig. S1) and cell type (Fig. 2), we found that MMD-MA, scJoint, LIGER and GLUE achieved a certain level of integration effect, and GLUE performed the best. MMD-MA could cluster cells well in scRNA-seq, while performed relatively worse in scATAC-seq.

**Fig. 2.**
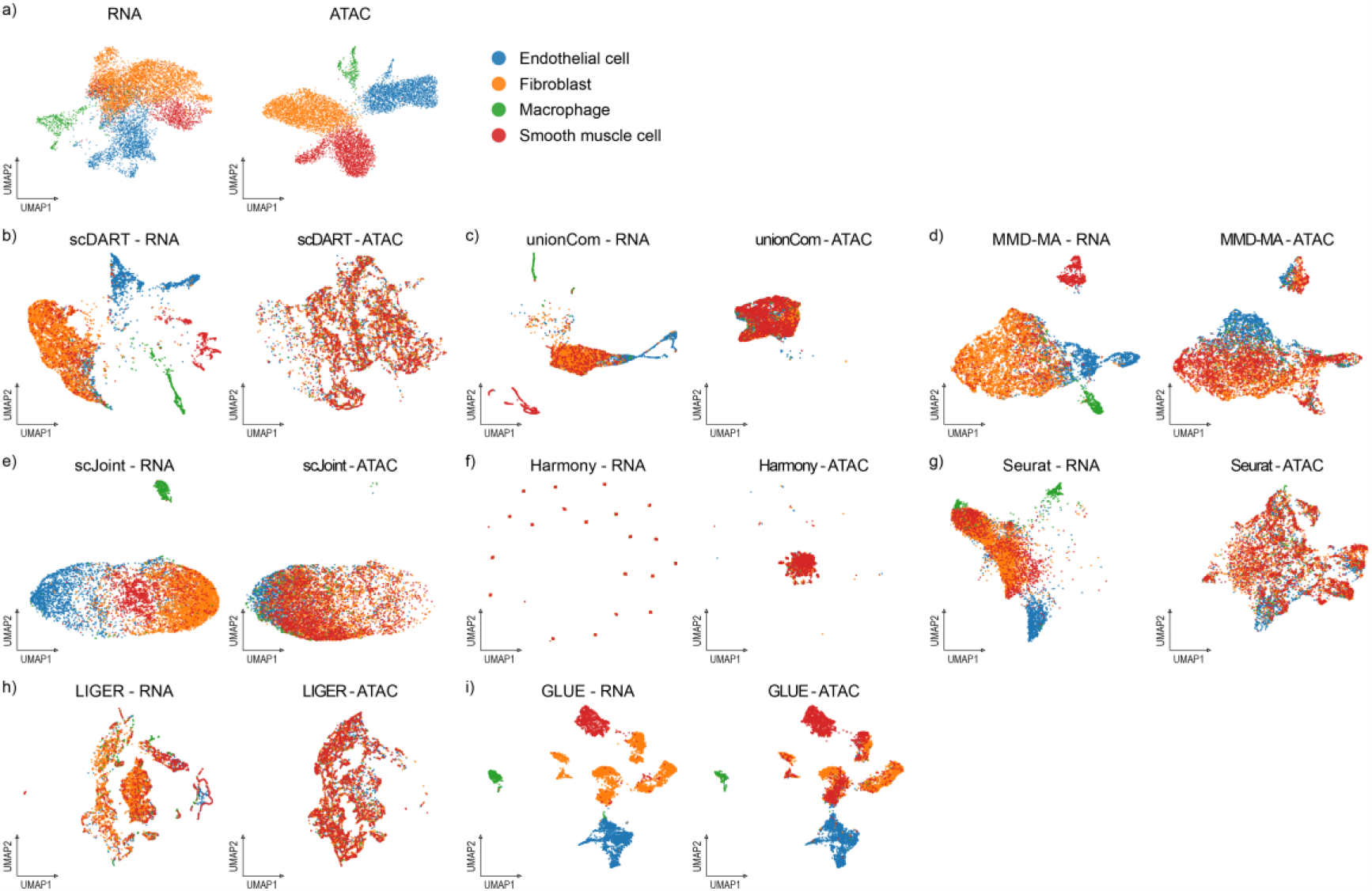
UMAP visualizations of the integrated cell embeddings for Dataset-U, colored by cell types. (a) Original visualization of cell types in two omics respectively. (b)-(i) Visualizations of the integration results of each method.

As shown Fig. 3, we found that GLUE had the highest scores in all metrics, and outperformed others greatly, especially in the score of NMI, showing its excellent performance in integrating the unpaired dataset. Meanwhile, MMD-MA, LIGER, Seurat had a relatively good score in omics mixing, and UnionCom, scJoint, scDART performed well in cell type conservation, indicating that they could also have a nice performance in one aspect.

**Fig. 3.**
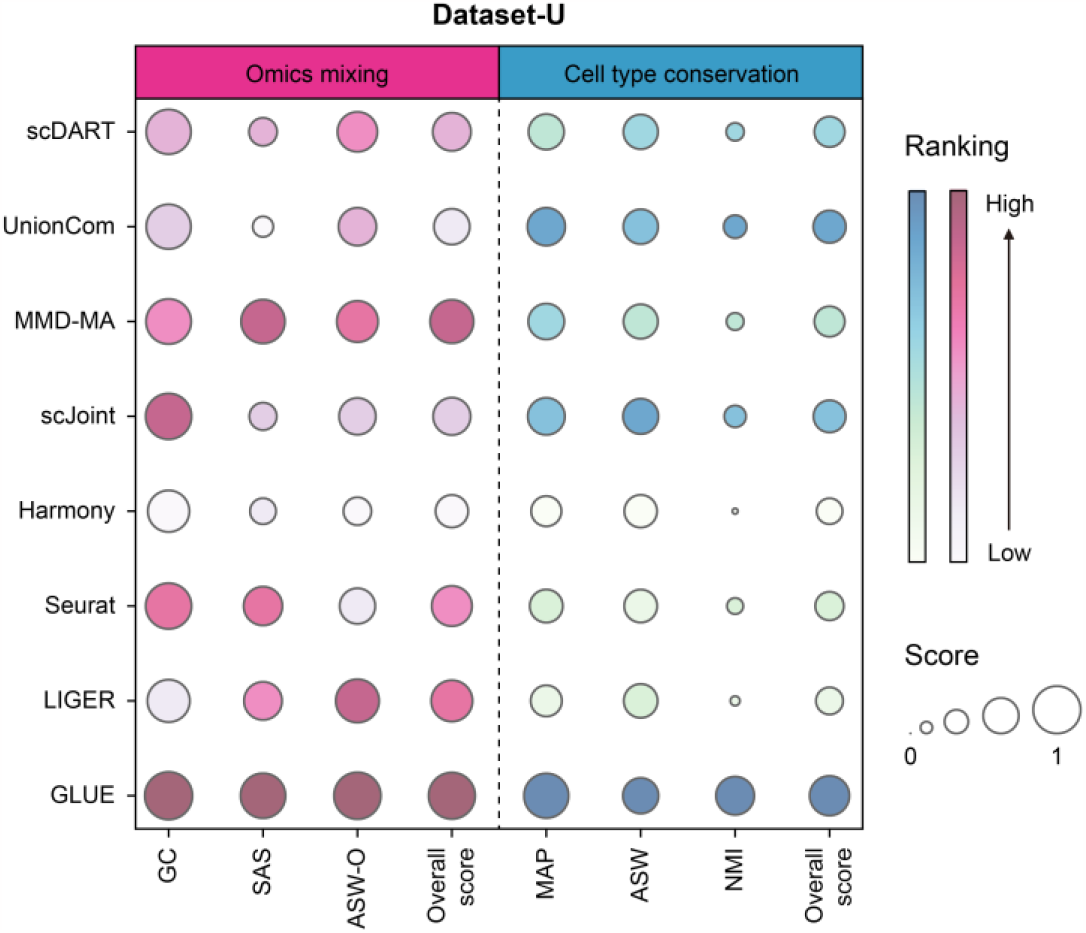
Detailed performance of evaluated methods on Dataset-U. Metrics of different categories were separated by dotted lines. The sizes of bubbles represented the scores of metrics, and the color represented the ranking.

### Benchmarking results on the paired dataset without a trajectory

Next, we wanted to evaluate how all the three kinds of methods perform in the task of integrating the paired dataset (Dataset-P). The 19 cell types could be divided from each other in scRNA-seq, but not clearly in scATAC-seq (Fig. 4a). In the UMAP visualizations that colored by omics type (Fig. S2), we observed that scDART, scJoint, LIGER, MultiVI and GLUE could mix two omics well, while UnionCom and Harmony performed relatively worse in this task. In the UMAP visualizations that colored by cell type (Fig. 4), we observed that scMVP, Seurat, MOFA+, MultiVI and GLUE achieved the best clustering effect in this task from visualization. scJoint and LIGER only achieved certain clustering effect in scRNA-seq, while MMD-MA clustered cell types inconsistently in two omics.

**Fig. 4.**
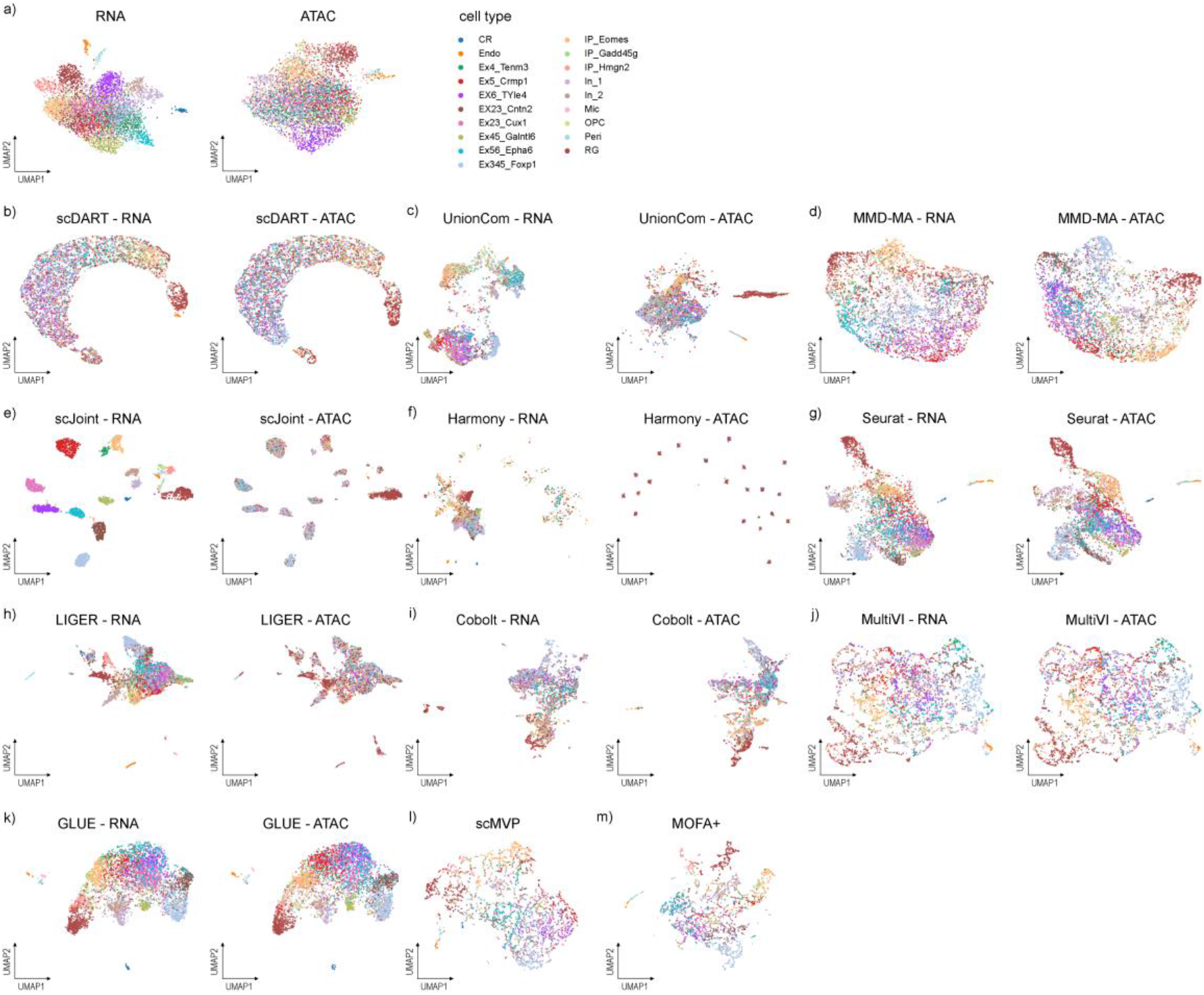
UMAP visualizations of the integrated cell embedding for Dataset-P, colored by cell types. (a) Original visualization of cell types in two omics respectively. (b)-(m) Visualizations of the integration results of each method. We visualized the distributions of cells in scRNA-seq and scATAC-seq separately for unpaired integration methods, and visualized the distribution in a single figure for paired integration methods.

We further calculated integration accuracy metrics to evaluate the performance of each method (Fig. 5). Since scMVP and MOFA+ didn’t outputted two omics in the latent space separately, we didn’t calculate their metrics in terms of omics mixing and alignment accuracy. Among all methods, GLUE outperformed others significantly in most metrics, especially in terms of cell type conservation and alignment accuracy, followed by MultiVI. Most methods except GLUE and MultiVI performed bad in NOS, which was used to measure whether two omics mix well when there was correspondence information, indicating that they may fail to integrate paired data one-to-one well. Besides, Seurat and LIGER performed well in omics mixing, and scMVP, MOFA+ performed well in cell type conservation. Although the performance of scJoint was not the best in each category, it achieved a relatively good balance among various aspects.

**Fig. 5.**
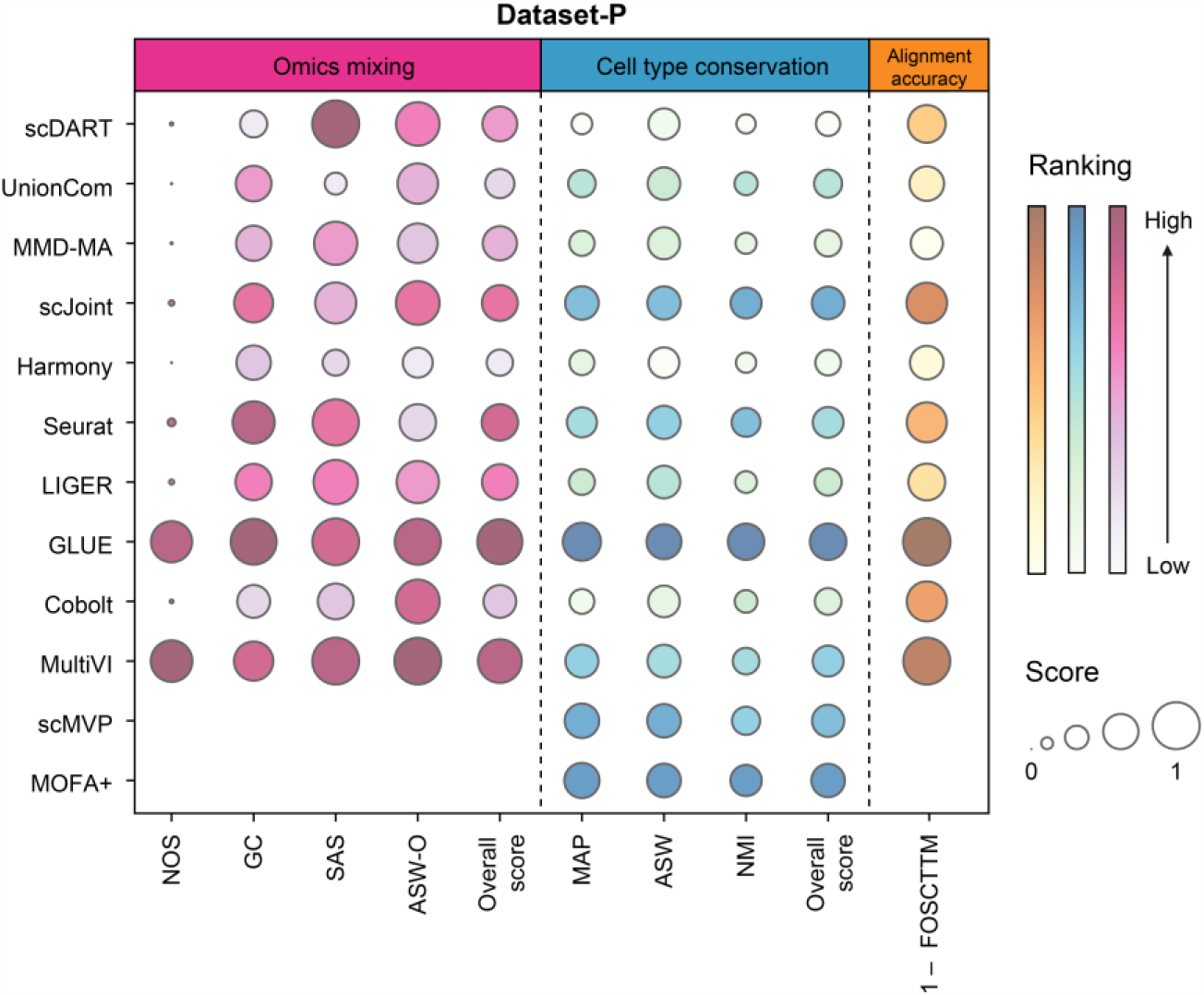
Detailed performance of evaluated methods on Dataset-P. Metrics of different categories were separated by dotted lines. The sizes of bubbles represented the scores of metrics, and the color represented the ranking. We used (1 – FOCSTTM) to represent the alignment accuracy to make the figure intuitive. As scMVP and MOFA+ didn’t output two omics in the latent space separately, we didn’t calculate their metrics in terms of omics mixing and alignment accuracy.

### Benchmarking results on the paired dataset with a trajectory

We evaluated how these methods performed in the task of integrating the paired dataset with a trajectory (Dataset-T). There was an expected linear trajectory going through IP-Hmg2, IP-Gadd45g, IP-Eomes, Ex23-Cntn and Ex23-Cux1 (Fig. 6a) [15]. From the UMAP visualizations that colored by omics type (Fig. S3), we found that scDART, MultiVI and GLUE could mix two omics well, while UnionCom, scJoint and Harmony performed relatively worse in this task. From the UMAP visualizations that colored by cell type (Fig. S4), we saw that scDART, Cobolt, scMVP and GLUE achieved the best clustering effects in this task, while scJoint and Seurat could only achieve clustering effects in the RNA omics.

**Fig. 6.**
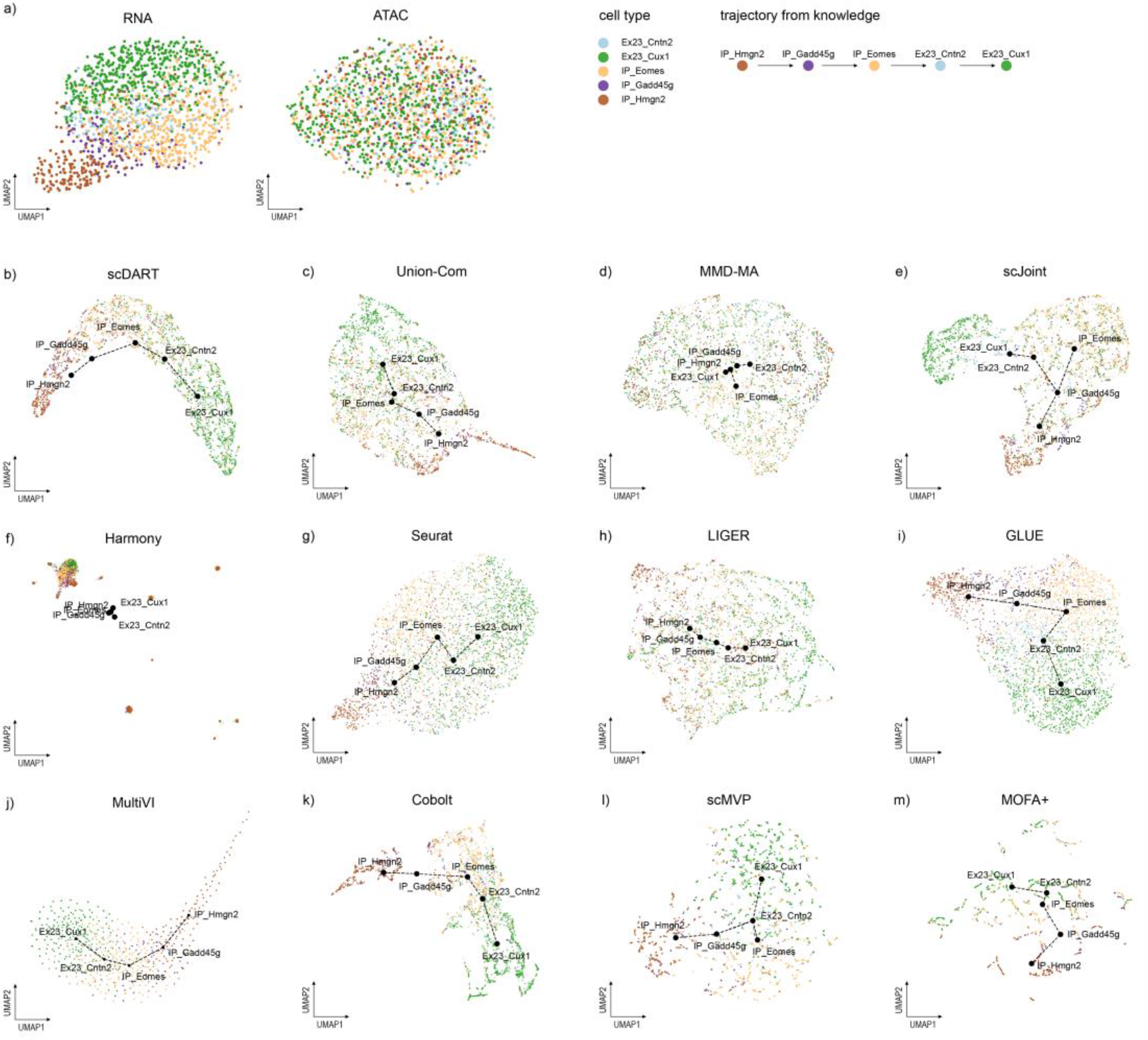
UMAP visualizations of the integrated cell embeddings for Dataset-T with trajectories, colored by omics types. Orange stands for scRNA-seq, and green stands for scATAC-seq. (a) Original visualizations of cell types in two omics respectively. (b)-(m) Visualizations of the integration results of each method, where the centroid of cells in each cell type were taken, and MST was used to obtain the trajectory on these points.

We then focused on the conservation of trajectories, and drew the UMAP visualizations with trajectories (Fig. 6). Specifically, we took the centroid of cells in each cell type, and used the Minimum Spanning Tree (MST) to obtain the trajectory on these points (Methods) [15]. We observed that most methods could have a good performance, while MMD-MA, scJoint, scMVP and Harmony didn’t preserve a linear trajectory, indicating their relatively poor function in this aspect.

Detailed integration accuracy metrics were shown in Fig. 7. Similar to the performance in Dataset-P, GLUE outperformed others in most metrics, and it was the only method who got a good score in NOS on this dataset. Besides, LIGER, scDART, MMD-MA had a good performance in omics mixing, and MultiVI, scMVP, Cobolt, MOFA+ performed well in cell type conservation. Only GLUE and MultiVI performed well in terms of alignment accuracy. As for the conservation of trajectories (Fig. 7), MultiVI, scMVP, MOFA+, GLUE had a good performance in F1 score, and MultiVI, GLUE, scDART performed well in correlation scores. After considering F1 score and correlation score comprehensively, MultiVI had the best performance in trajectory conservation, followed by GLUE.

**Fig. 7.**
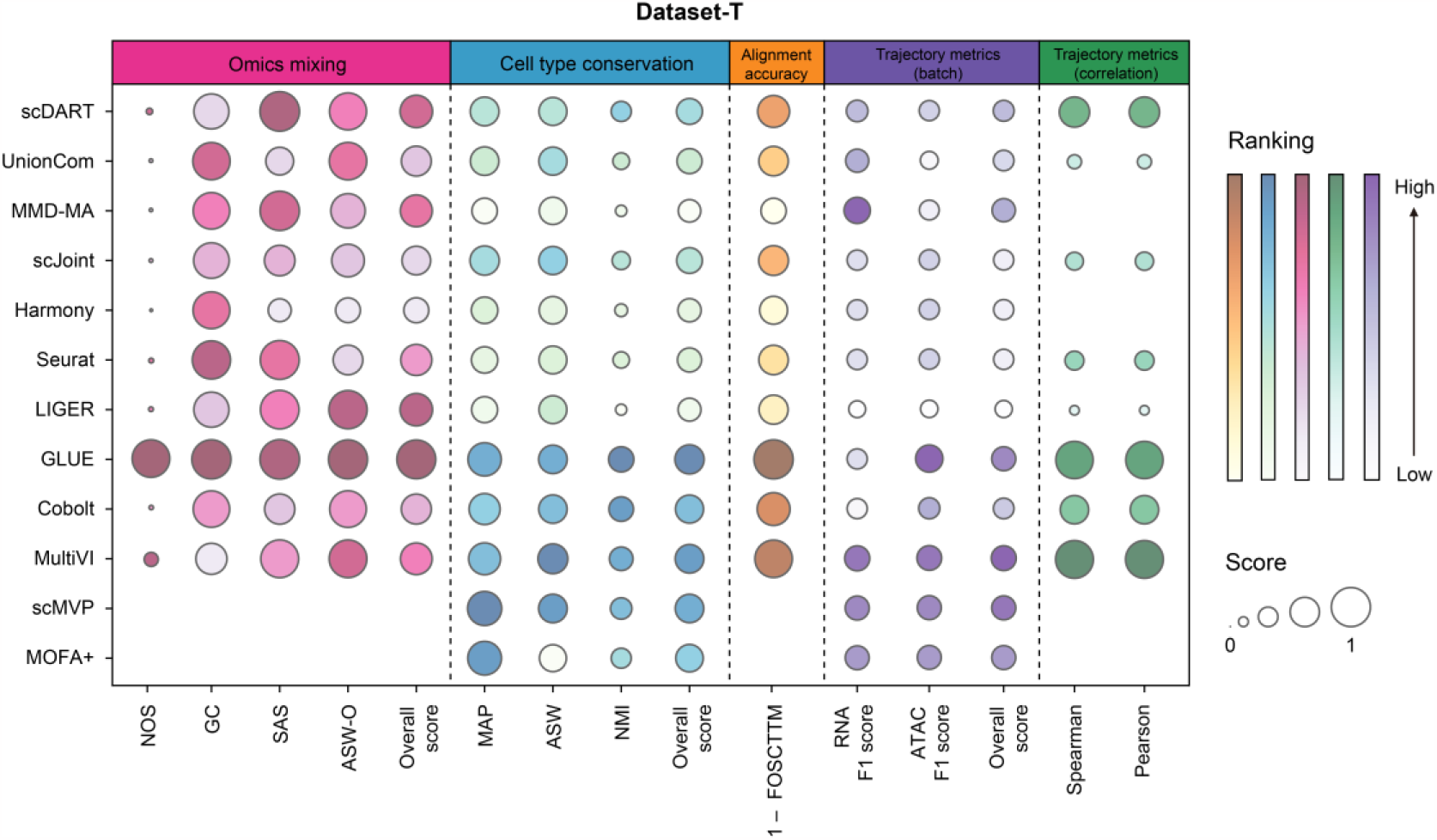
Detailed performance of evaluated methods on Dataset-T. Metrics of different categories were separated by dotted lines. The sizes of bubbles represented the scores of metrics, and the color represented the ranking. We used (1 – FOCSTTM) to represent the alignment accuracy to make the figure intuitive. As scMVP and MOFA+ didn’t outputted two omics in the latent space separately, we didn’t calculate their metrics in terms of omics mixing, alignment accuracy, and correlation score. As the scores of Spearman and Pearson correlation for MMD-MA and Harmony were negative, the results were not shown in the bubble chart.

### Scalability and ease of use

In order to compare the scalability of each method, we monitored the running time when testing each dataset (Fig. 8a). As expected, integrating more cells led to longer runtime. We found that scJoint, Harmony, Seurat, LIGER and MOFA+ performed best in terms of runtime, and the runtime of scDART, UnionCom and MMD-MA did increase obviously with the dataset size. The runtime of GLUE didn’t show a significant increasing trend when the number of cells increased, but it had the longest time to integrate multi-omics when there were only over a thousand cells.

**Fig. 8.**
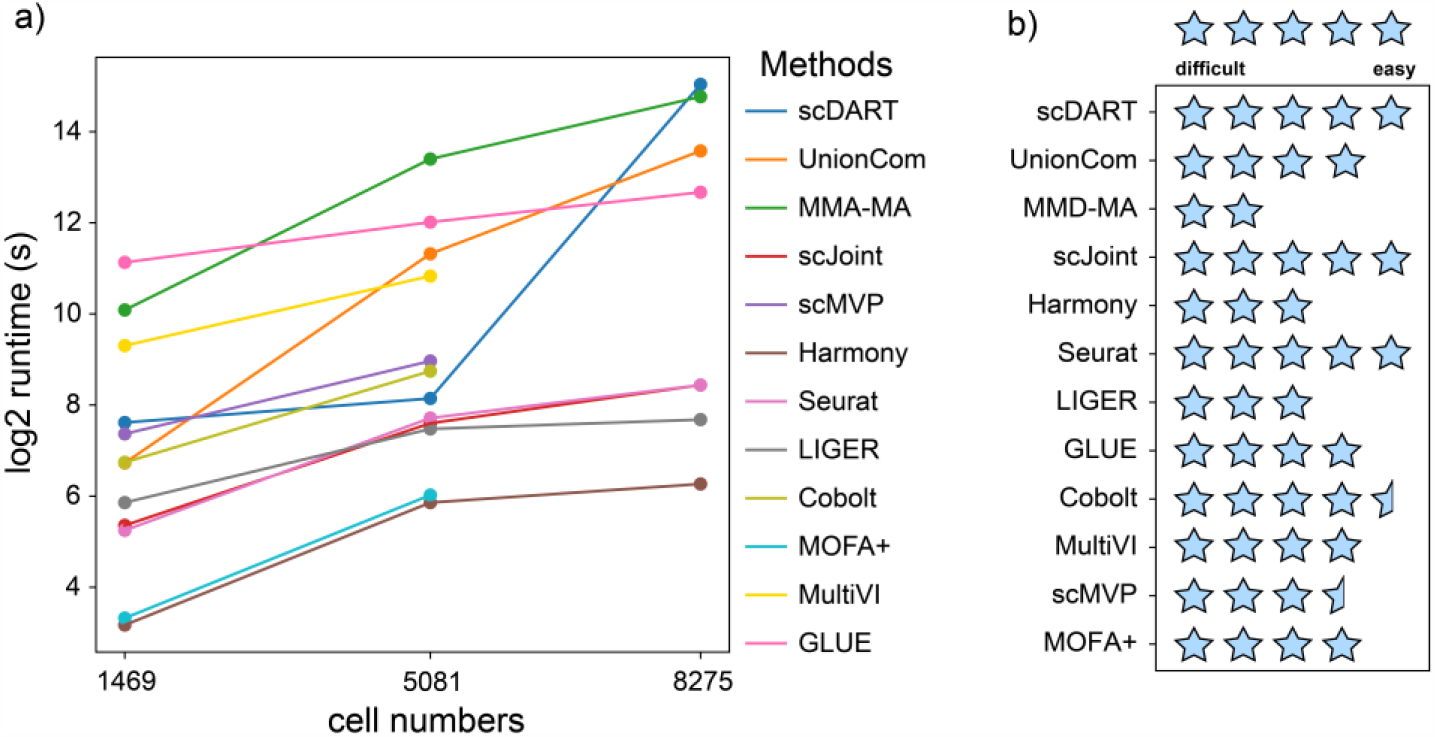
(a) Scalability (running time) for each method. Methods that can only be used for paired datasets were not tested for the unpaired one (cell number = 8275). (b) Ease of use for each method.

During the testing approach, we found that these methods showed varied degrees of ease to use. we thus graded the ease of use for each method based on our experience (Fig. 8b). This grading may be somewhat subjective can help the users, especially researchers not specialized in bioinformatics, choose the suitable method. We first investigated the guidance provided by these methods. Some methods provided detailed web pages for guiding users, such as Seurat (Integrating scRNA-seq and scATAC-seq data · Seurat (satijalab.org)), MOFA+ (Tutorials | Multi-Omics Factor Analysis (biofam.github.io)), MultiVI (MultiVI — scvi-tools), GLUE (https://scglue.readthedocs.io/zh-cn/latest/). Some methods have detailed tutorial files in their GitHub pages, including scDART, scJoint, scMVP, Harmony, LIGER, Cobolt. Such detailed guidance could make it easier for users to do experiment on their own datasets. We also considered our experience in testing these methods. Some methods are relatively easy to implement, such as Seurat, UnionCom, Harmony, LIGER, scDART, and scJoint. The deep learning-based methods are generally more complicated than traditional methods for usage. Considering the two aspects together, we suggested Seurat, scDART and scJoint as the easiest methods to use.

## Discussion

We benchmarked 12 integration methods on 3 integration tasks via 8 metrics that measured their performance between omics mixing, cell type conservation and alignment accuracy. We used 3 additional metrics to evaluate the preservation of trajectory on the dataset with a trajectory. Besides, we monitored the running time when testing each dataset to analyze the trend of their runtime as the number of cells increased and graded these methods based on their ease of use. We drew radar plots to comprehensively compare the performance of different aspects for each method (Fig. 9), which could help users have a more intuitive understanding and thereby helping them make better choices among these methods.

**Fig. 9.**
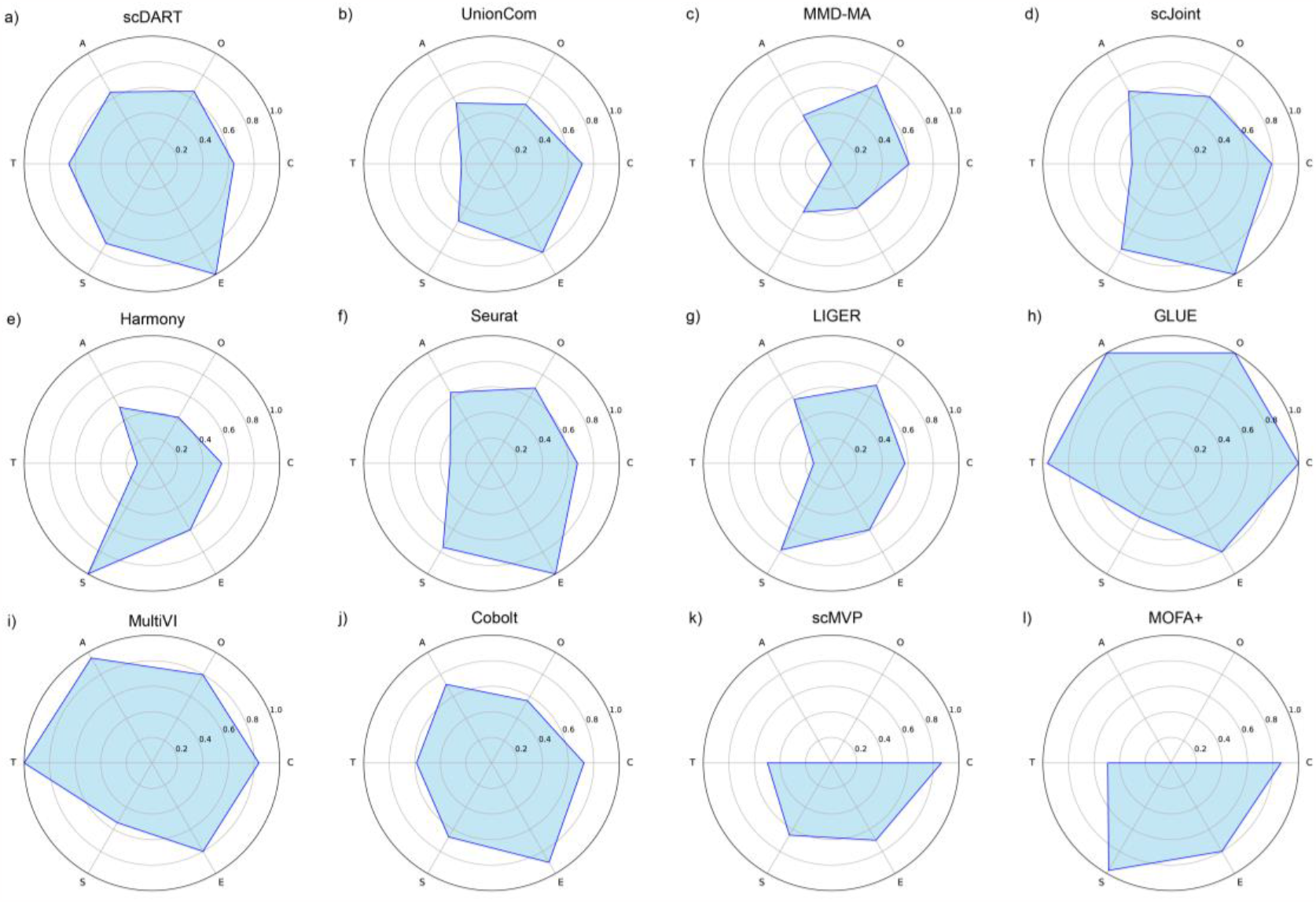
Performance of all benchmarked methods on different aspects. The abbreviations in the figure represented: O (omics mixing), C (cell type conservation), A (alignment accuracy), T (trajectory conservation), S (scalability), E (ease of use). We calculated the average score in every aspect for each method, and scaled the scores by dividing the maximum value of this aspect. For scalability, we chose the runtime that cell number = 8275, took the reciprocals of the log2-transformed values, and scaled them by dividing the maximum value. For paired methods, their scores of omics mixing and alignment accuracy were set to 0.

For the unpaired dataset, GLUE was the top performer. Whether from visualization or metrics perspective, its performance far surpassed other methods, indicating its excellent performance in integrating unpaired datasets. Besides, MMD-MA, LIGER, Seurat performed relatively well in omics mixing, and UnionCom, scJoint, scDART also had relatively good scores in cell type conservation.

When dealing with paired datasets, most methods could achieve a good integration effect especially when the number of cells was small. But with the number of cells and cell types increasing, some methods failed to maintain its functions. Specifically, GLUE outperformed others in terms of omics mixing, cell type conservation and alignment accuracy, followed by MultiVI. Besides, in terms of omics mixing, LIGER, Seurat and scDART could also mix two omics well. MOFA+, scMVP performed well in terms of cell type conservation. In addition to the above methods, Cobolt also performed well when the number of cells was not big.

As for trajectory conservation, MultiVI, scMVP, MOFA+, GLUE and scDART had good scores in certain metrics, and MultiVI performed the best after considering multiple metrics comprehensively, followed by GLUE.

After considering multiple factors comprehensively, we would advise to choose the integration method according to the category and size of datasets. For example, when integrating unpaired datasets, we recommend to choose GLUE. As for paired tasks, it may up to integration purposes and the scale of datasets. Generally, GLUE would also be the best choice, followed by MultiVI. And these two methods are also the best choices for trajectory conservation. In addition, if one focuses on omics mixing, scDART, LIGER, and Seurat are also worth a try. As for cell type conservation, MOFA+, scMVP could also be taken into consideration. Considering scalability, Seurat, LIGER and MOFA+ may save running time. Besides, scDART, scJoint and Seurat can be used with an easier start with the detailed guidance they provided.

Overall, we hope this work could help researchers to better evaluate the performance of these popular integration methods in the field of single-cell RNA and ATAC data integration, and could help them to have a better thinking of how to choose methods. Additionally, we hope that the benchmarking strategies can benefit the future development of novel multi-omics integration methods.

## Methods

### Testing environment

During testing, Harmony (v.0.1.1), LIGER (v.1.0.0) and Seurat (v.4.3.0) were based on R language (4.2.0). scMVP was based on Python 3.7.12. MultiVI was based on Python 3.9.16. GLUE was based on Python 3.10.0 and package ‘scglue’ (v.0.3.2). Cobolt was based on Python 3.8.16. MOFA+ was based on Python 3.8.0. Other methods were based on Python language (3.10.8). The hardware environment used during the test was GeForce RTX 2080 Ti, and the CPU model was 72 Intel (R) Xeon (R) Gold 5220 CPU @ 2.20GHz, with 512GB of memory.

### Datasets

#### The unpaired dataset (Dataset-U)

These two human uterus datasets were got from different work. Specifically, scRNA-seq was generated from Wang et al. [25], and scATAC-seq was generated from Zhang et al. [26]. We selected four cell types that were common in both scRNA-seq and scATAC-seq. Next, we randomly filtered the cell types with the highest number of scRNA-seq and scATAC-seq in each of the two omics, ensuring that the order of cell numbers for each type was consistent in both omics. This resulted in 8,237 cells for scRNA-seq and 8,314 cells for scATAC-seq.

#### The paired dataset without a trajectory (Dataset-P)

This dataset was a P0 mouse cerebral cortex dataset with 5,081 cells generated by droplet-based SNARE-seq, downloaded from MCBI GEO accession number GSE126074, with both raw gene expression and DNA accessibility measurements available for the same cell [1]. The scRNA-seq data had 19,322 gene features, while the scATAC-seq had 229,429 peak features, and the cells could be divided into 19 cell types. The cell-type information was obtained from the previous study [20].

#### The paired dataset with a trajectory (Dataset-T)

This dataset was extracted from the dataset mentioned above, which included only 1469 cells and 5 cell types. These five cell types (IP-Hmgn2, IP-Gadd45g, IP-Eomes, Ex-L2/3-Cntn2 and Ex-L2/3-Cux1) measured the differentiation trajectory from intermediate progenitor cells to upper-layer excitatory neurons [1,15]. Further, features were selected for both omics, resulting in 933 gene features for scRNA-seq and 15,857 peak features for scATAC-seq. The detailed data and cell-type information were obtained from the previous study [15].

#### Data preprocessing and settings used in each method

For Dataset-U, we tested eight methods: scDART, UnionCom, MMD-MA, scJoint, Harmony, Seurat, LIGER and GLUE. For two paired datasets, we additionally tested four methods: scMVP, Cobolt, MOFA+ and MultiVI. Detailed data preprocessing and settings for each method were shown as follows, which were according to the treatments mentioned in their papers. We also ran these methods with and without scaling and HVG selection if there were no corresponding instructions in the papers, and chose the settings corresponding to the best result for each method after multiple attempts.

- scDART: Region2gene matrix was generated by the code provided by the original paper [15]. For Dataset-P, the dimension of latent space was 4, and trained for 500 iterations. And we selected top 1,000 most variable genes in scRNA-seq using Scanpy, and the corresponding 24,286 peaks in scATAC-seq using the code provided by the original paper [15]. For Dataset-T, the settings were the same except that we didn’t select HVGs. For Dataset-U, we set the dimension of latent space as 4 and batch size as 4, and trained for 500 iterations. We also selected the top 1,000 most variable genes in scRNA-seq using Scanpy, and the corresponding 32,524 peaks in scATAC-seq.
- UnionCom: For Dataset-P, the dimension of latent space was 32, and trained for 2,000 iterations. We normalized and log-transformed data after selecting top 1,000 most variable genes in scRNA-seq, and binarized and reduced the dimensionality of scATAC-seq to 1,000. For Dataset-T, the dimension of latent space was 32, and trained for 10,000 iterations. The following data preprocessing was the same as above, except that we didn’t select HVG for scRNA-seq. For Dataset-U, the only difference with the settings for Dataset-P was that we trained for 10,000 iterations.
- MMD-MA: For Dataset-P, we set *l*_1_ as 1e-5, *l*_2_ as 1e-5, the dimension of latent space as 5, bandwidth as 0.5, seed as 50, and trained for 10,000 iterations. We normalized and log-transformed data after selecting top 1,000 most variable genes in scRNA-seq, and reduced the dimensionality of scRNA-seq to 100. For scATAC-seq, we binarized the features and did TF-IDF transformation, and reduced the dimensionality to 100. For Dataset-T, the only difference was that we directly reduced the dimensionality of scRNA-seq to 100. For Dataset-U, all the settings were the same as Dataset-P.
- scJoint: For Dataset-P, we set batch as 256, *l*_*r*1_ and *l*_*r*3_ as 0.01, epoch_1_ and epoch_3_ as 50, center_weight as 0.1, crossentropy as TRUE, and the dimension of latent space as 64. We binarized both omics as inputs. For Dataset-T, the difference was that we set epoch_1_ and epoch_3_ as 25, and center_weight as 50. For Dataset-U, we set center_weight as 1.
- scMVP: For the two paired datasets, we pretrained scRNA-seq and scATAC-seq following the code it provided. For Dataset-P, we set the dimension of latent space as 10, and for Dataset-T, we set the dimension of latent space as 20.
- Harmony: For Dataset-P, we set *λ* as 0.1, plot_convergence as TRUE, the max iterations as 20, and the dimension of latent space as 50. For Dataset-T, the difference was that we set the max iterations as 50. The settings for Dataset-U were the same as Dataset-T.
- Seurat: For three datasets, the settings were all default, with 50 as the dimension of the latent space.
- LIGER: For Dataset-P, we set the dimension of latent space as 19, *λ* as 5, thresh as 1e-6, and the max iterations as 50. We selected top 1,000 most variable genes in scRNA-seq and corresponding peaks in scATAC-seq as we did in scDART mentioned before. For Dataset-T, we set the dimension of latent space as 5, the max iterations as 30, and didn’t select HVGs. For Dataset-U, the settings were the same as Dataset-P except that we set the dimension of the latent space as 4.
- Cobolt: For Dataset-P, we set lr as 0.001, the dimension of latent space as 30, and trained for 100 iterations. For both omics, we did log-plus-one transformation, and applied quality filtering as the code it provided [23]. Then the data was divided into 20% as paired and 80% as unpaired, following the tutorial. For Dataset-T, we set lr as 0.005, the dimension of latent space as 10, also trained for 100 iterations. The following preprocessing was the same as above, except that we didn’t do quality filtering.
- MOFA+: For Dataset-P, we set the dimension of latent space (*k*) as 10, and did log-transformed normalization for both omics. Then we selected top 2,500 most variable genes for scRNA-seq and top 5000 most variable genes for scATAC-seq. For Dataset-T, the settings were the same, except that we didn’t select HVGs for scRNA-seq.
- MultiVI: For Dataset-P, we set seed as 420, batch_key as *modality*, lr as 1e-4, and max iterations as 500. The data was divided into 20% as paired and 80% as unpaired, following the tutorial [22]. For Dataset-T, the only difference was that we set lr as 5e-5.
- GLUE: For all three datasets, we did the preprocessing just like the tutorial [17], and set the dimension of latent space as 50, and use_obs_names as TRUE.

#### Visualization

Inspired by the code of Zhang et al. [15], we implemented three plotting functions in python. For Dataset-P and Dataset-U, we drew the UMAP visualizations of the integrated cell embeddings colored by omics types and cell types separately for each method. For Dataset-T with a trajectory, we additionally drew a UMAP plot with a trajectory for each method. The latent space data of two omics were first superimposed, and then the centroids of each cell type were taken to construct a Minimum Spanning Tree (MST), which represents the reservation of trajectory.

#### Evaluation Metrics

Inspired by previous studies [15,17,27], we selected some metrics that were useful in determining the integration effect. These metrics could be divided into four categories: omics mixing, cell type conservation, alignment accuracy at single-cell level, and trajectory conservation. The first category contained neighborhood overlap score (NOS), graph connectivity (GC), Seurat alignment score (SAS) and the average silhouette width across omics (ASW-O). Scores from the second category included mean average precision (MAP), the average silhouette width (ASW) and normalized mutual information (NMI). Alignment accuracy was evaluated via FOSCTTM. As for the last category, metrics included F1 score across branches (F1) and Spearman and Pearson correlation.

#### Omics mixing

*neighborhood overlap score (NOS)*. NOS could be used to evaluate the degree of mixing between omics when there existed cell-cell correspondence across data modalities. First, the *k*-nearest neighbor of scRNA-seq and scATAC-seq were calculated respectively in the latent space, and then the proportion of matching cells corresponding to this cell in the nearest neighbor of the other omics was evaluated. This metric required prior information on the correspondence between data in two omics, which could only be applied in the case of paired integration. The obtained value was between 0 and 1, and the higher the value, the better the omics mixing effects.

*graph connectivity (GC)*. GC could be used to evaluate the mixing between omics, and its calculation was defined as [17,27]:

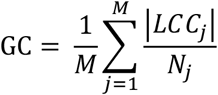

Where *LCC*_*j*_ denoted the number of cells with the largest connected component in the k-nearest neighbor graph for cell type *j, N*_*j*_ was the number of cells in cell type *j*, and *M* was the total number of cell types. The obtained value was between 0 and 1, and the higher the value, the better the omics mixing effect.

*Seurat alignment score (SAS)*. SAS could also be used to evaluate the mixing between omics, and its calculation was defined as [17,28]:

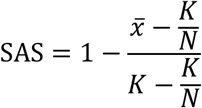

Where 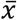 was the average number of cells from the same omics layer in the *k*-nearest neighbors of cells (different layers were first sampled to the same number of cells as the smallest layer), and *N* was the number of omics layers. The obtained value was between 0 and 1, and the higher the value, the better the omics mixing effect.

*average silhouette width across omics (ASW-O)*. ASW-O could also be used to evaluate the mixing between omics, and its calculation was defined as [17,27]:

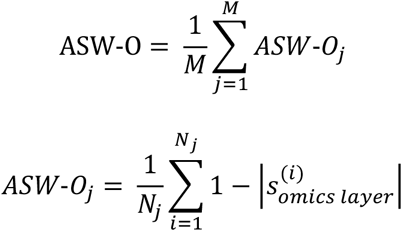

Where 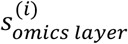 was the omics layer silhouette width for the *i*th cell, *N* _*j*_ was the number of cells in cell type *j, M* was the total number of cell types. The obtained value was between 0 and 1, and the higher the value, the better the omics mixing effect.

*Omics mixing*. Based on the above four metrics, we could calculate the final omics mixing score following the procedure of previous studies [17,27]:

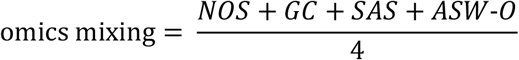

#### Cell type conservation

*mean average precision (MAP)*. MAP could be used to evaluate the cell type resolution. Supposing that the cell type of the *i*th cell was y^(*i*)^, and the first *K* nearest neighbors were 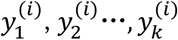 then MAP was calculated as [17]:

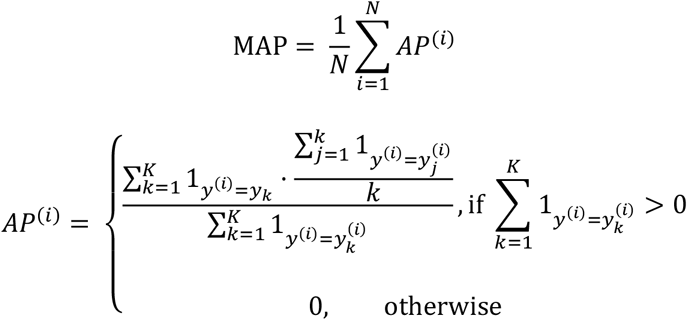

Where 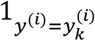 was the indicator function, which equaled to 1 when 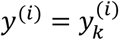 and 0 otherwise. For each cell, AP calculated the average cell type accuracy of the neighbors matched for each cell type, while MAP was the average AP value for all cells. The obtained value was between 0 and 1, and the higher the value, the better the cell type resolution.

*average silhouette width (ASW)*. ASW could also be used to evaluate the cell type resolution. Its calculation was defined as [17,27]:

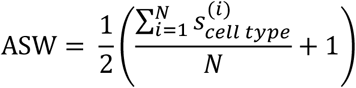

Where 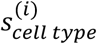 was the cell type silhouette width of the *i*th cell. The obtained value was between 0 and 1, and the higher the value, the better the cell type resolution.

*normalized mutual information (NMI)*. NMI could be used to measure the overlap of two clusters [27]. We first performed Louvain clustering to obtain the best match between clusters and labels. And then used NMI to compare the cell type label with Louvain clusters computed on the integrated results. The overlap score was calculated using the mean of the entropy terms for cell type and cluster labels. The obtained value was between 0 and 1, and the higher the value, the better the cell type resolution.

*Cell type conservation*. Based on the above three metrics, we could calculate the final cell type conservation score following the procedure of previous studies [17,27].

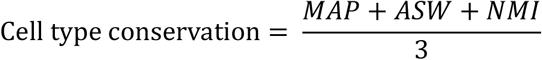

#### Alignment accuracy

*FOSCTTM*. FOSCTTM was used to evaluate the alignment accuracy at single-cell level. Assuming that each omics data contains *N* cells, and these cells were sorted in the same order in both omics, that is, the *i*th cell in omics one was paired with the *i*th cell in omics two. Let *x* and *y* represent the cell coordinates of omics one and omics two in latent space respectively, then FOSCTTM was calculated as [17,30]:

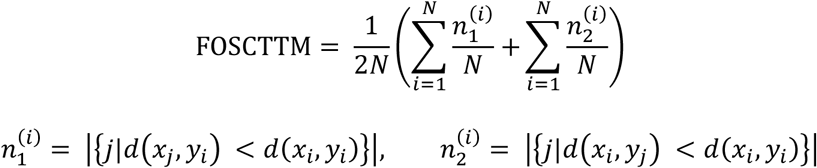

Where 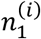 and 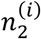 were the number of cells closer to the original paired cell in the other omics. This metric required prior information on the correspondence between data in two omics, which could only be applied in the case of paired integration. The obtained value was between 0 and 1, and a smaller value indicated a higher alignment accuracy.

#### Trajectory conservation

*F1 score (F1)*. F1 score across branches could be used to evaluate the accuracy of cell branch assignment, which was an aspect for the accuracy of trajectory [15,29]. Given the distribution of the ground truth and the inferred cell branches, we first calculated the Jaccard similarity between each pair of inferred cell branches and the ground truth branches. And we calculated the *recovery* and the *relevance* as the average maximum Jaccard similarity for every branch in ground truth and in inferred branches respectively. The F1 score was then defined as [15,29]:

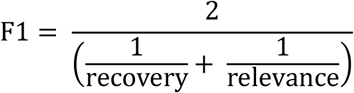

The obtained value was between 0 and 1, and a higher value indicated a better assignment for cell branches.

*Spearman and Pearson correlation*. Because matched cells should ideally have the same pseudotime along trajectory, these two metrics could be used to evaluate the consistency of pseudotime inferred from cells in both omics [15]. We first inferred the pseudotime of each omics on the latent space respectively, and then calculated the correlation of pseudotime between omics using Spearman and Pearson correlation. The obtained value was between –1 and 1, and a higher value indicated a better consistency between pseudotime.

## Funding

The work is supported in part by National Key R&D Program of China (grant 2021YFF1200900), and National Natural Science Foundation of China (grants 62250005, 61721003, 62373210).

## Author contributions

C.X., Y.C., L.W. and X.Z. conceived the study. L.W. and X.Z. supervised the research. C.X. and Y.C. designed and implemented the experiments. C.X., Y.C. and L.W. prepared the figures. C.X., Y.C., L.W. and X.Z. wrote the manuscript, C.X., Y.C., L.W. and X.Z. reviewed the manuscript.

## Data availability

The SNARE-seq dataset (Dataset-P and Dataset-T) can be downloaded from the GEO website (https://www.ncbi.nlm.nih.gov/geo/), with number GSE126074. For scRNA-seq in Dataset-U, related data are available at the GEO website GSE111976. For scATAC-seq in Dataset-U, related data are available at the GEO website, with number GSE184462.

## Code availability

The source codes for tests and evaluations are available online on GitHub at https://github.com/sunnsset/Benchmarking_single-cell_RNA_and_ATAC_data_integration.

## Declaration of interests

The authors declare no competing interests.

## Supplementary Figures

**Fig. S1.**
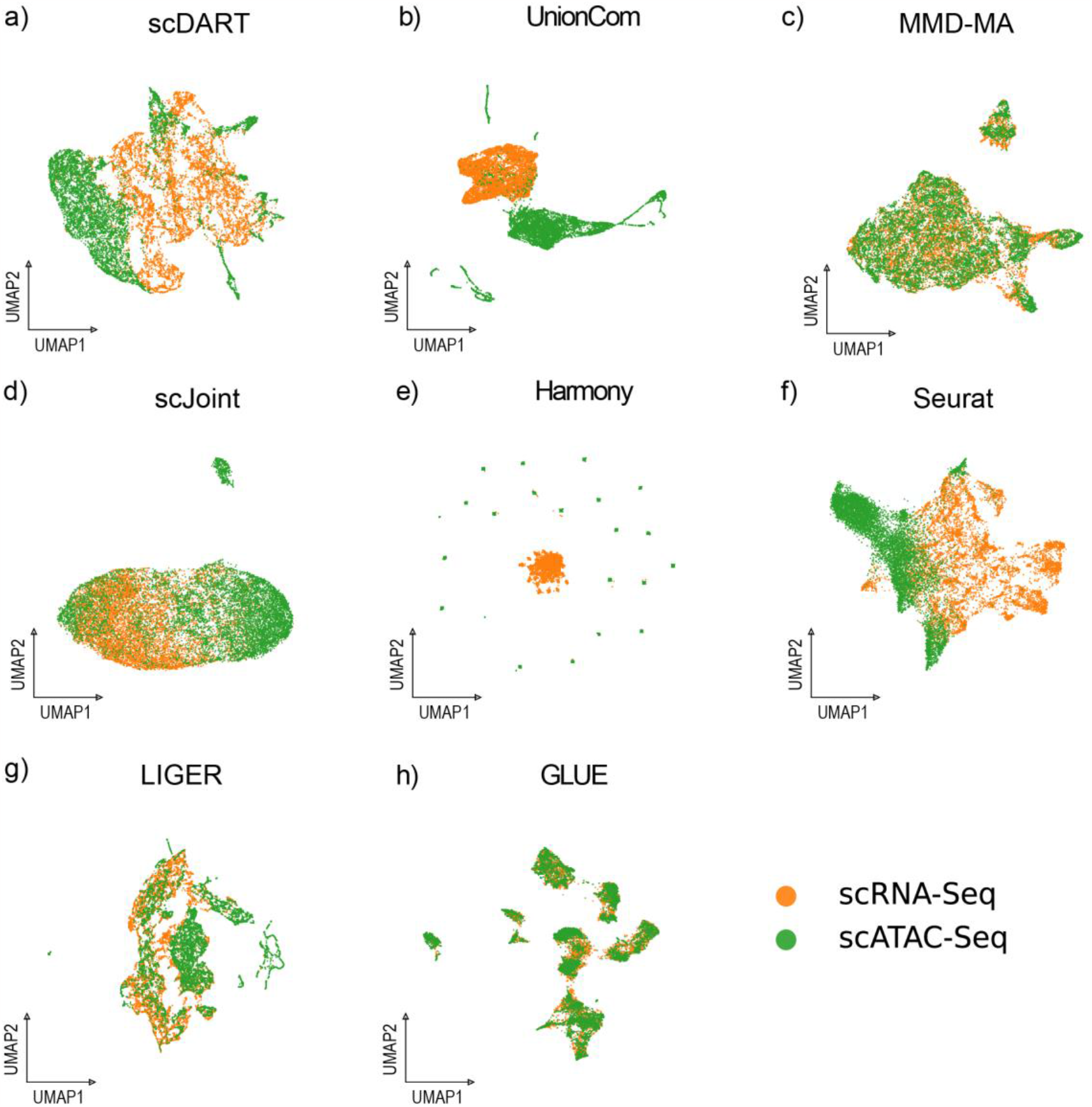
UMAP visualizations of the integrated cell embedding for Dataset-U, colored by omics types. Orange stands for scRNA-seq, and green stands for scATAC-seq.

**Fig. S2.**
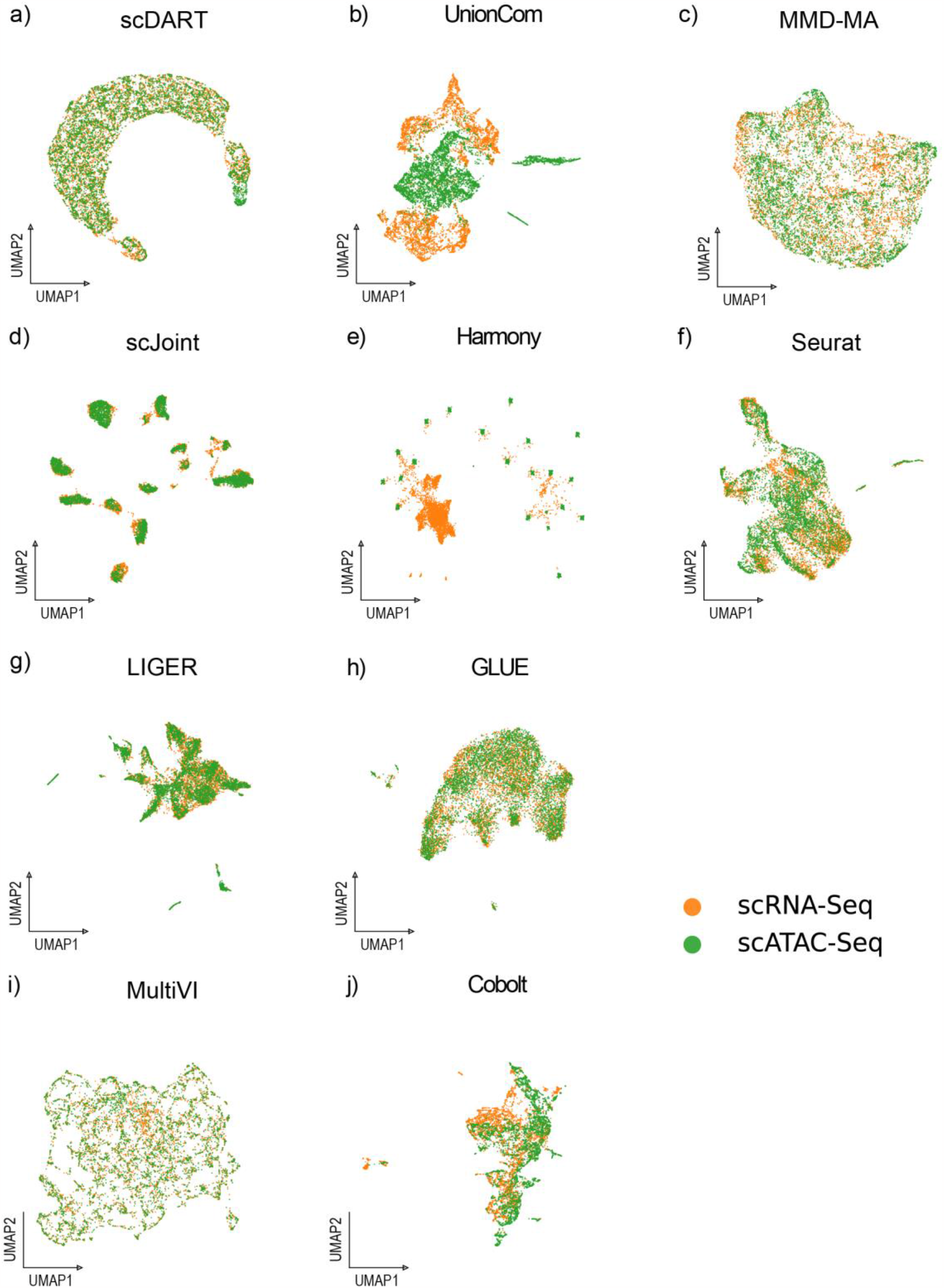
UMAP visualizations of the integrated cell embedding for Dataset-P, colored by omics types. Orange stands for scRNA-seq, and green stands for scATAC-seq. As the latent distributions of two omics were not separately provided by scMVP and MOFA+, this diagram for these two methods were not shown here.

**Fig. S3.**
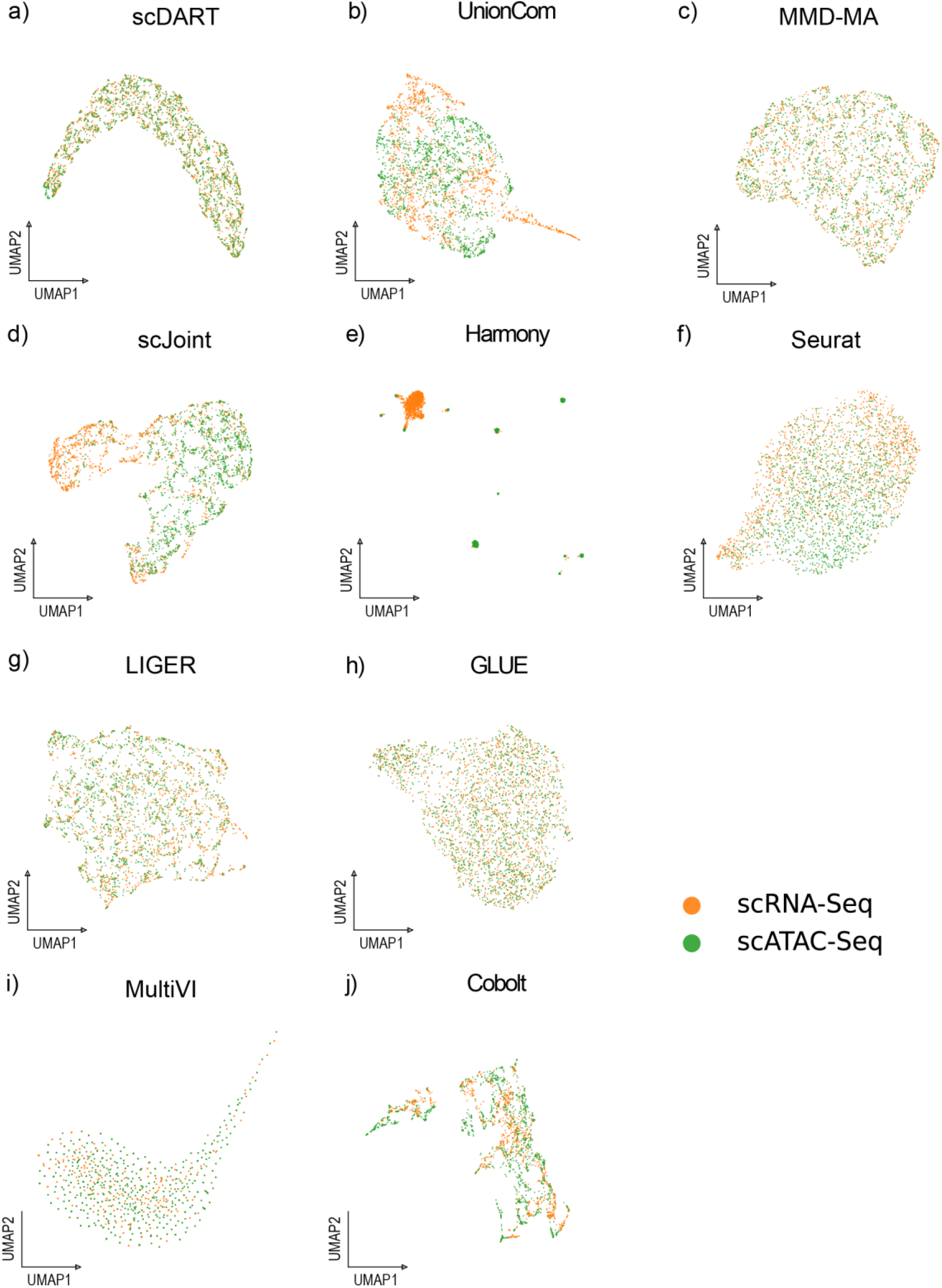
UMAP visualizations of the integrated cell embedding for Dataset-T, colored by omics types. Orange stands for scRNA-seq, and green stands for scATAC-seq. As the latent distributions of two omics were not separately provided by scMVP and MOFA+, this diagram for these two methods were not shown here.

**Fig. S4.**
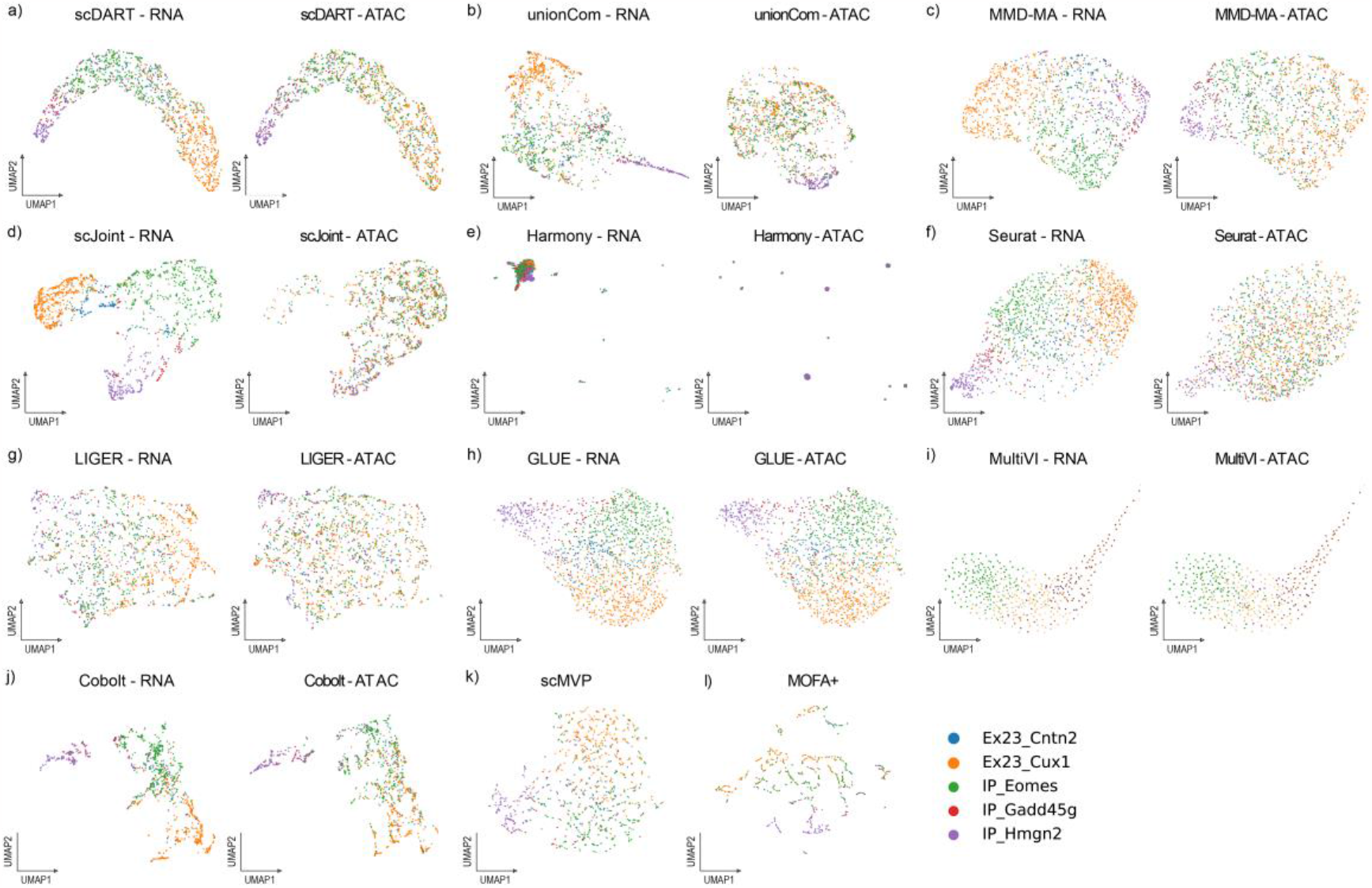
UMAP visualizations of the integrated cell embedding for Dataset-T, colored by cell types. We visualized the distributions of cells in scRNA-seq and scATAC-seq separately for unpaired integration methods, and visualized the distribution in a single figure for paired integration methods.

## Notes

### Competing Interest Statement

The authors have declared no competing interest.

